# Influence of truthful and misleading instructions on statistical learning across the autism spectrum

**DOI:** 10.1101/2024.07.12.603256

**Authors:** Nikitas Angeletos Chrysaitis, Peggy Seriès

## Abstract

Bayesian studies of perception have documented how the brain learns the statistics of a new environment and uses them to interpret sensory information. Impairments in this process have been hypothesised to be central to autism spectrum disorders. However, very few such studies have differentiated between implicit and explicit learning. We manipulated the instructions given before a cue-stimulus association task to investigate their effects on statistical learning. The task was conducted online, in 335 participants with varying autistic traits. In the implicit condition, where no information was provided, participants acquired weak prior beliefs about the task regularities. Conversely, explicit information about the presence of regularities resulted in strong priors, correctly reflecting the task’s statistics, regardless of the information’s veracity. Contrary to our hypothesis, autistic traits did not correlate with the influence of priors in any condition.

**Author Summary:** Perception is greatly influenced by the brain’s prior knowledge of the environment, through a process called *Bayesian inference*. Recent theories of psychiatric disorders and particularly autism view them as impairments in this process. A crucial aspect of this process is how individuals form their knowledge of the environment. However, previous studies have not differentiated between learning that occurs when participants are aware of what they are learning and learning that happens implicitly. In the present study, we conducted an experiment with four conditions, each varying in terms of what participants were trying to learn and whether they were aware of the general form of the regularities. Our findings revealed that participants form much stronger beliefs about the regularities when they are informed about their presence. Additionally, we discovered that participants with strong autistic traits did not differ in their beliefs about the regularities in any of the conditions.

## Introduction

The information we get from our senses is frequently noisy and ambiguous. To overcome that, the brain combines sensory evidence with prior knowledge about the environment. This process of integrating sensory inputs with prior beliefs is formalised in Bayesian theories of perception and cognition [1]. According to Bayesian models, the brain maintains internal, probabilistic models of the environment. These are combined with sensory information that takes the role of the likelihood in Bayesian inference to generate posterior probability distributions over possible states of the world. The posteriors are then used to form what we end up perceiving and how we understand the world.

There are different ways that the brain could learn about the world. It could be that gradual experience with the regularities of the environment results in implicit prior beliefs. Alternatively, knowledge of the regularities can be explicitly communicated to the individual. This raises a few questions. Do priors that are based on explicit information differ from those that are formed implicitly? Does misleading information affect subsequent learning? And does that differ when there are no regularities in the environment versus when there are, but they differ from what you were told?

In this study, we set to answer these questions. Previous research on implicit and explicit learning has mostly focused on serial reaction time tasks and artificial grammar learning [2,3]. The results show evidence for two distinct processes [4,5], affecting different brain areas [6– 8]. These processes depend on different factors, with explicit learning being correlated with IQ [9,10] and the resulting knowledge decaying over time [11,12], while implicit learning is largely independent from intelligence and its effects appear to be consolidated with the passage of time. Interestingly, when they are compared on the same task, alerting participants to the presence of regularities often leads to inhibition of implicit learning and worse participant performance [7,13–15]. However, to our knowledge, no studies have investigated implicit and explicit learning when environmental regularities are based on the frequency of the stimuli, instead of specific rules that they follow. Accordingly, no studies have investigated participant behaviour from a Bayesian viewpoint.

Our study is partially inspired by the Bayesian literature on autism spectrum disorder (ASD). Initially, such theories proposed that autistic individuals exhibit weaker prior influences compared to the general population [16–18]. While this remains the main hypothesis, in recent years the focus has shifted towards explanations that are centred on the learning of regularities and specifically the processing of environmental volatility [19,20]. Indeed, our systematic review found that ASD groups differed more often from neurotypical groups in tasks where priors were learned compared to tasks involving pre-existing priors [21]. Our review also showed that studies using implicitly learned priors tended to find weaker influences in autistic individuals, while this was less common when prior learning was explicit or when participants were directly informed about the regularities [21]. This seems to align with non-Bayesian studies showing intact explicit but occasionally impaired implicit learning in ASD [see 22 for a review]. Unfortunately, most ASD studies did not describe in detail what instructions were given to the participants, meaning it was sometimes unclear what components of the learning were implicit or explicit. Moreover, the two kinds of tasks commonly used very different experimental designs, making the comparison between them difficult.

In the present study, we used one design, manipulating the regularities and the instructions of the task independently across four conditions to assess their individual impact on learning and their interaction. The participants were asked to discriminate between left- and right-tilted Gabor patches that were preceded by an auditory cue. The cue-stimulus relationship varied across conditions, as did the participants’ knowledge of it. We modelled the participant behaviour using a Bayesian model, so that we could estimate the participants’ priors and compare them between conditions. We also collected responses on the autism spectrum quotient questionnaire (AQ) and investigated the potential relationships between these scores and participant performance on the task. We found that when participants were not given any information about the regularities, they developed very weak beliefs about them. In contrast, when their attention was drawn to the presence of regularities, their beliefs were much stronger, independently of the veracity of the information they received. We found no relationship between participant behaviour and autistic traits in any task condition.

## Methods

### Participants

We recruited 456 participants from the crowdsourcing platform Prolific, with approximately half of them having reported that they have a diagnosis of ASD or that they feel they belong in the autism spectrum (Section 1 in S1 Supplementary Information). All participants completed the AQ questionnaire and then one of the conditions of our task. All participants gave informed consent and received monetary compensation for participation. The study was approved by the University of Edinburgh School of Informatics Research Ethics Committee (Application number 175134). As a quality control, we discarded the data of the participants who failed our attention checks, who did not respond in more than 3.5% of the trials, or who could not discriminate between high- and low-pitch tones. This was done before any behavioural or computational analysis. Surprisingly, more than one fourth of the participants were excluded via this process, leaving us with 335 participants across all conditions. The group that self-identified as part of the autism spectrum had fewer rejections than the other group (20% vs 32%). AQ scores did not differ between accepted and rejected participants (mean AQ 24.9 vs 24.4, *p* = 0.47).

### Design

The experiment consisted of a cue-stimulus association task (Figure 1). In all conditions, the experiment began with instructions explaining that the goal of the task was discriminating between left- and right-tilted (Gabor) gratings. Then participants completed one training block of 64 trials that included only the visual stimuli. The task began at a very easy level, with the gradings being presented at a very high contrast and at 70° or 120°. During the training, the contrast changed based on a 2/1 perceptual staircase (expected to yield 71% correct responses upon convergence), while the tilt of the stimuli was gradually reduced to 93° and 87°, respectively. This was done so that participants would get gradually acclimated to the task. The contrast staircase also led to stimuli that were similarly hard to see across our sample, as we could not control the environment of online participants. In each trial, gratings were presented for 150ms and were immediately followed by a static target consisting of four concentric circles for 300ms to prevent the formation of afterimages. Then, participants had 1000ms to choose if the grating was tilted to the left or to the right, using the left and right arrow keys. The screen displayed two arrows (oriented at 45° and 135°, see Figure 1B) at that time corresponding to the possible orientations. When participants made their choice, the other arrow disappeared and the remaining arrow turned green or red for 300ms, depending on if they were correct or not.

**Figure 1:**
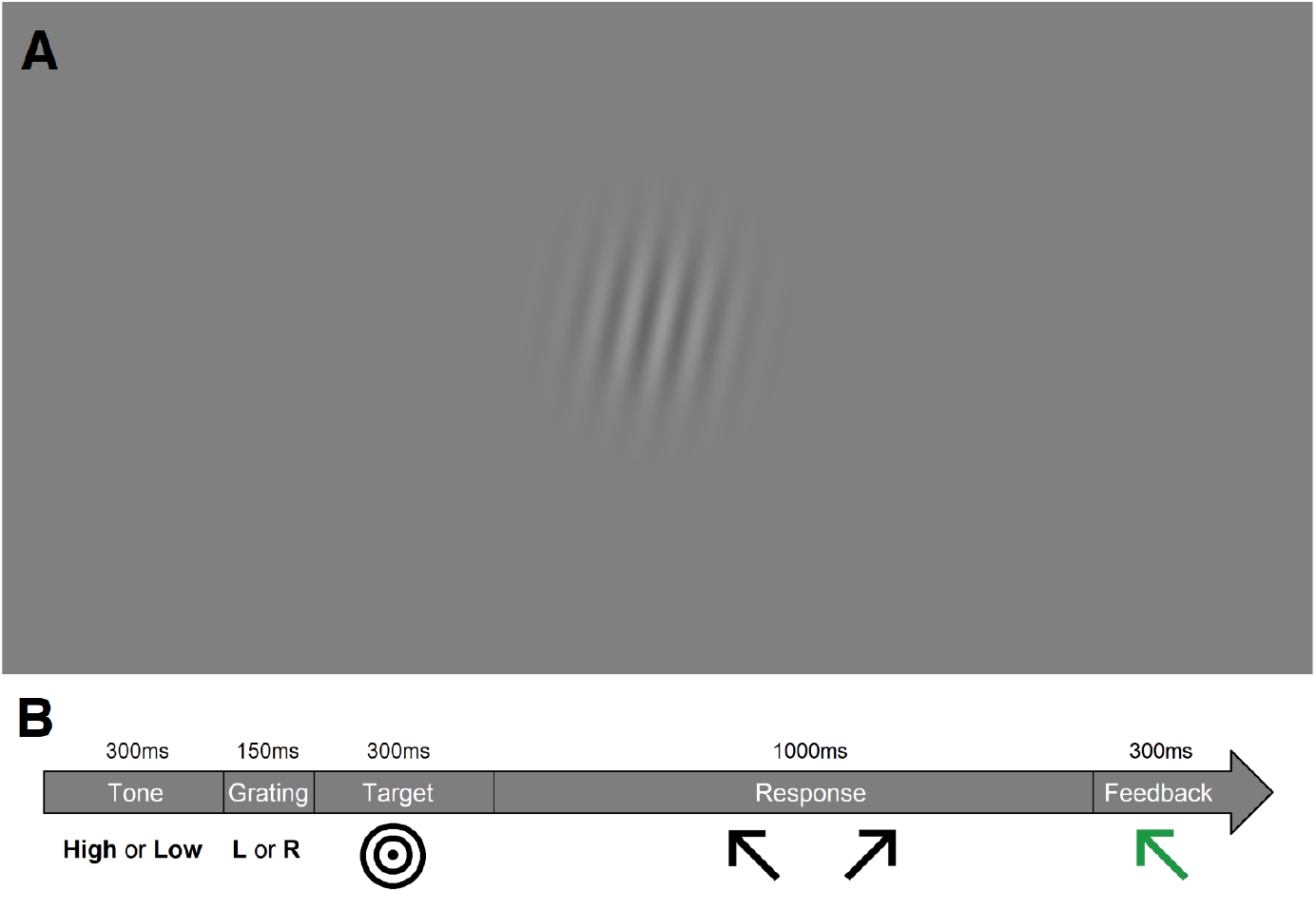
Example stimulus (A) and structure of individual trial (B). Stimuli in the experiment were presented in significantly lower contrast than the example in (A). After the stimulus, the screen showed a target followed by two arrows (shown in B: Response). After the participants responded the arrow they pressed flashed green if their response was correct or red if it was not (illustrated in B: Feedback).

The trials of the main task kept the same structure with two changes. Firstly, the contrast of the grating started at the value set by the training and was updated using a 3/1 staircase (expected to yield 79% correct responses upon convergence). Secondly, participants were initially presented with a random pure tone of either 330hz or 660hz for 300ms before the presentation of the grating. The relationship between the auditory cues and the visual stimuli differed across conditions, as did the instructions given to the participants about them. The main task lasted for 4 blocks or 256 trials.

Participants were split into four conditions based on the information they received from the instructions (I) and the regularities (R). Receiving no explicit information about the regularities or the tone not being associated with the image is denoted with the number 0. Otherwise, the presence of information is denoted with a + symbol, or with a – when instructions disagreed with the regularities.

- In the condition I^0^R^+^, the implicit condition, the tone predicted the tilt of the grating with 75% probability. The pairing of the tone to the tilt was randomised between participants. Participants were naïve about the association between tones and gratings (instructions: ‘The purpose of the tone is to alert you to the presence of the grating and focus your attention on it.’). 119 participants took part in this condition.
- In the condition I^+^R^0^, the tone had no relationship with the grating (50%). Participants, though, were told that ‘A low-pitch tone will be followed 75% of the time by a [right/left]-tilted grating. A high-pitch tone will be followed 75% of the time by a [left/right]-tilted grating’. Thus, participants were given explicit information about the regularities, but this information was misleading as the true regularities were completely random. 106 participants took part in this condition.
- The final two conditions were a combination of the above, with the tone predicting the tilt of the grating with 75% probability and the instructions being the same as in the second condition. In approximately half the participants (I^+^R^+^, n = 51) the instructions were telling the truth, similarly to explicit conditions in the literature. In the other half (I^−^R^+^, n = 59) the association mentioned in the instructions was the reverse to the true association. That is, if in truth the low tone was followed 75% of the time by a left-tilted grating, the instructions said that it was followed 75% of the time by a right-tilted one and vice versa. The purpose of these conditions was to see how explicit information interacted with the learning of the regularities, both when agreeing and when disagreeing. The I^−^R^+^ condition has some similarities with the I^+^R^0^ condition as in both the instructions were misleading. However, it is possible that learning the regularities of an environment employs different mechanisms from learning that there are no regularities or learning to simply ignore the instructions.

Participants were similar across conditions in the distribution of their AQ scores, their age, and their gender (Section 1 in S1 Supplementary Information). We expected that in all conditions but the implicit one (I^0^R^+^), the instructions would create a prior belief in the participants that would bias their responses at the beginning of the experiment. In the I^+^R^0^ condition, participants would gradually learn that the information provided was false and they would end up with no biases. In the rest, participants would gradually learn the actual regularities. This would result in the I^+^R^+^ condition forming the strongest priors by the end of the experiment, followed by the I^0^R^+^ condition. Prior development in the I^−^R^+^ condition would be hindered by the instructions but would eventually update towards the regularities. We also hypothesised that participants with high AQ scores would develop weaker priors in the implicit, I^0^R^+^ condition, while they would show no differences in the I^+^R^+^ condition, as they would be aided by the instructions.

### Computational Modelling

Besides simple statistical methods, we analysed the behaviour of the participants using four computational models. One was a simple response model, which assumed that participants responded in the task either based on the auditory or on the visual stimulus but did not combine the information from the two. Mathematically,

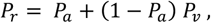

where *Pr* is the probability of responding ‘right’ in a single trial, *Pa* is the probability of simply following the auditory cue, and

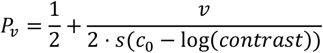

with v = 1 if the stimulus is tilted to the right and v = –1 if it is tilted to the left. *c*_0_ is a participant-specific parameter that represents a baseline contrast level and *s* is the standard logistic function.

Our second model was a static Bayesian model which was formalised as

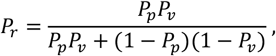

where *Pp* is the prior based on the auditory cue and *Pv* functions as the likelihood function. Our third model followed the same core Bayesian structure, but had a dynamically changing *Pp*, modelled by the Hierarchical Gaussian Filter [HGF, 23]. The HGF is a Bayesian model which tracks the evolution of beliefs over time as well as the volatility of the environment. The final model, was a simpler learning model that also used the same core Bayesian structure but modelled the evolution of participant beliefs using a Rescorla-Wagner model. The mathematical details for both models can be found in Section 3 in S1 Supplementary Information The four models were fit on all trials and compared using Bayesian model averaging [24].

## Results

### Behavioural Analysis

We expected the prior to bias participants’ responses towards the directions that were associated with the tones. This would result in higher accuracy when tilts and tones were congruent and lower when they were not (Figure 2). By ‘congruent’, we refer to trials where the tone-tilt pairings correspond to the most likely pairing (appearing in 75% of the trials, in the I^0^R^+^, I^+^R^+^, and I^−^R^+^ conditions) or to the pairing participants were told would appear in 75% of the trials (in the I^+^R^0^ condition). Note that in the I^−^R^+^ condition we are defining ‘congruent’ according to the actual regularities and not the instructions. Specifically, we expected that in the I^0^R^+^ condition participants would start with no difference between congruent and incongruent trials and would gradually develop a bias during the experiment. On the other hand, in the I^+^R^0^ condition, participants would start being strongly biased and then they would realise that there are no regularities and gradually reduce or even eliminate their bias. Then, the I^+^R^+^ condition would be a combination of the I^0^R^+^ and I^+^R^0^ ones. Participants would start biased and would remain at the same level throughout the experiment. Finally, in the I^−^R^+^ condition, they would start being biased in the opposite direction and then would gradually develop a bias in the correct direction, albeit never reaching the magnitude of the bias of the I^0^R^+^ or I^+^R^+^ conditions.

**Figure 2:**
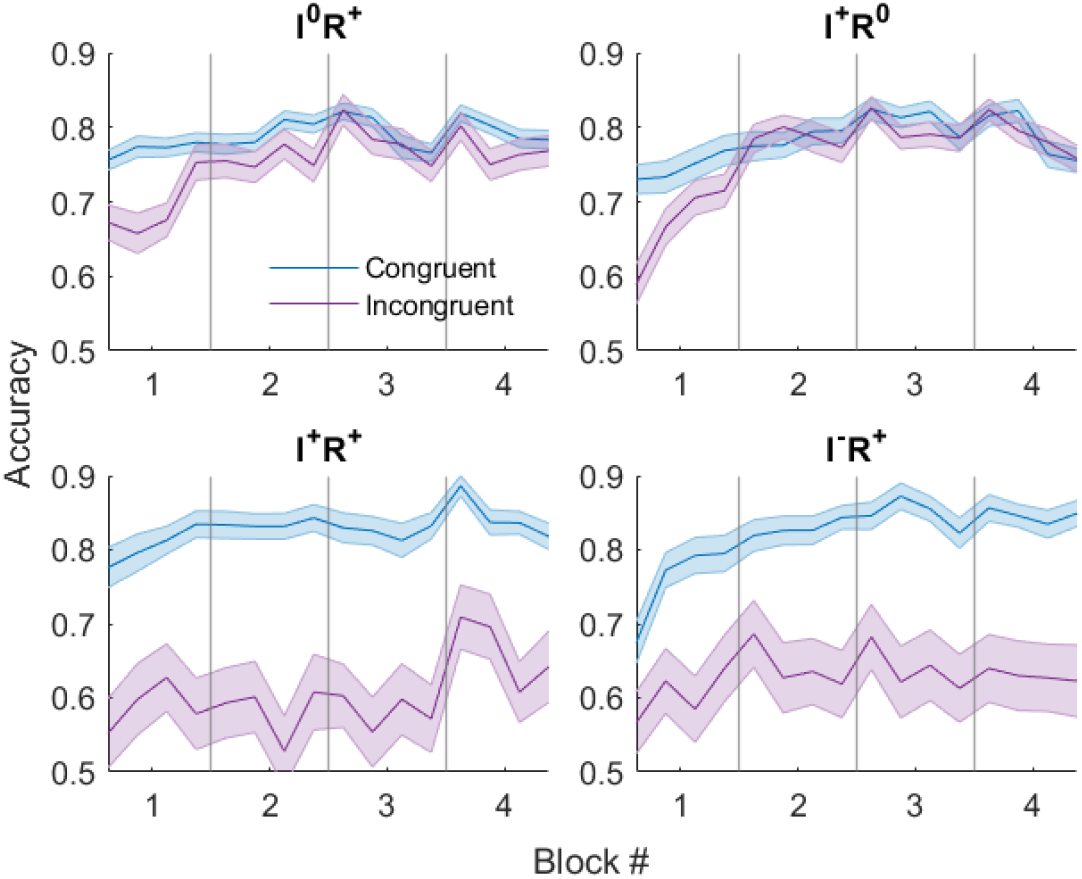
Accuracy comparison between congruent and incongruent trials in the four conditions. Participants showed small accuracy differences in the implicit condition, large accuracy differences in the two conditions where explicit information was combined with regularities, and no differences in the I^+^R^0^ condition after the first block. Each datapoint corresponds to one fourth of a block or 16 trials. Blocks are separated by vertical lines. Coloured areas correspond to standard errors.

Looking at the accuracy differences (Figure 2), we see that participants showed biases in the direction of the actual environmental regularities. A two-way mixed ANOVA yielded a significant effect of congruency (F = 118.9, *p* = 10^−23^), which differed significantly across conditions (F = 30.3, *p* = 10^−17^). Post-hoc t-tests showed that participants had significantly higher accuracy in the congruent trials than in the incongruent ones in all conditions besides I^+^R^0^ (Table 1). Biases in the I^+^R^+^ and I^−^R^+^ conditions were larger than those in the I^0^R^+^ condition (*t* = 7.59, *p* = 10^−12^, BF_10_ = 10^9^ and *t* = 5.78, = 10^−8^, BF_10_ = 10^5^, respectively), but they did not differ among each other (*t* = 0.39, *p* = 0.69, BF_10_ = 0.22). Reaction times were slightly smaller in the congruent condition (F = 4.1, *p* = 0.045), but this explained approximately 1% of response variance (η^2^ = 0.01) and was not different across conditions (F = 0.5, *p =* 0.70).

**Table 1.** Accuracy differences between congruent and incongruent trials for blocks 2-4.

| Condition | Bias | $t$ | $p$ | $BF_{10}$ |
| --- | --- | --- | --- | --- |
| $I^0R^+$ | 0.024 | 2.7 | 0.007 | 3.7 |
| $I^+R^0$ | 0.004 | 0.5 | 0.65 | 0.12 |
| $I^+R^+$ | 0.226 | 6.5 | $10^{-8}$ | $10^5$ |
| $I^-R^+$ | 0.204 | 5.0 | $10^{-6}$ | $10^3$ |

From Figures 2 and 3, it appears that participants were able to learn the regularities within the first block, at least partially, and they did not update their beliefs in the following trials. This was verified by fitting the participant biases to the block numbers using a linear mixed-effects model, which yielded no difference across blocks 2, 3, and 4 (F = 0.1, *p* = 0.78, BF_10_ = 10^−7^) and no interaction between blocks and conditions (F = 1.0, *p* = 0.38, BF_10_ = 10^−5^). However, as the staircase which determined the contrast changed between the main experiment and the training, it could be that these results are influenced by the contrast slightly increasing during the experiment (Section 2 in S1 Supplementary Information). When the contrast is lower, the participants have to rely on their priors more and any potential biases are stronger. This could also explain the unexpected trajectory of the bias in the I^0^R^+^ condition, which starts relatively high and then drops within the first block: participants developed a weak prior quickly, whose influence was reduced as the contrast increased. Attempting to account for the effects of contrast we analysed the data using computational modelling.

**Figure 3:**
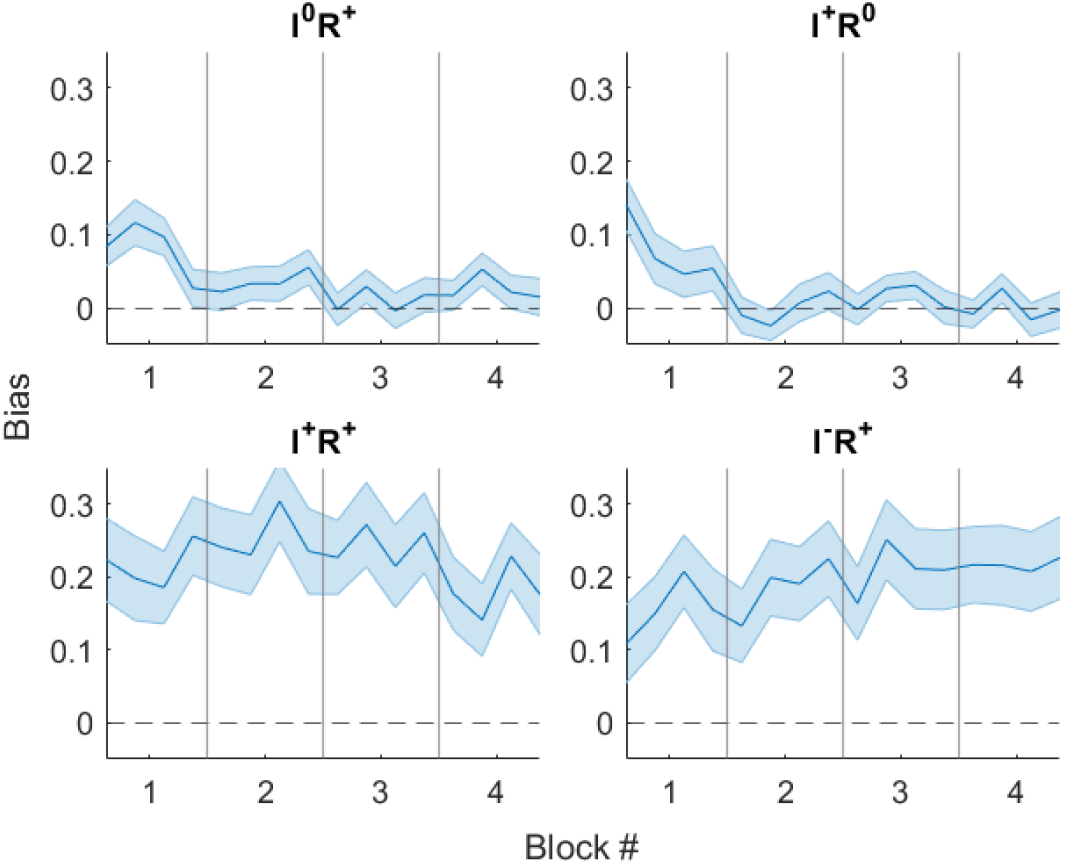
The evolution of bias over time in the four conditions. Each datapoint corresponds to one fourth of a block or 16 trials. Blocks are separated by vertical lines. Coloured areas correspond to standard errors.

### Computational Analysis

Comparing different models on all trials (Section 4 in S1 Supplementary Information), we found that the best model in all conditions was the static Bayesian one, despite not being able to model the acquisition of the participants’ priors. This could potentially be explained by the fact that participants developed priors very quickly, as evident by the exhibition of biases since the first trials of the first block, which then stayed stable throughout the task. In contrast with that, the estimated trajectories of participant priors by the HGF appear to increase throughout the task (Figure 2 in S1 Supplementary Information), perhaps reflecting the inability of the HGF to model this type of learning.

As the static Bayesian model cannot simulate participant learning, we made a post-hoc decision to refit the static Bayesian model only on trials from the second, third, and forth blocks before analysing its behaviour. The estimated priors for the four conditions can be seen on Figure 4. Mirroring the behavioural results, priors were significantly different across conditions (F = 30.9, *p* = 10^−17^, BF_10_ = 10^14^), with priors in all conditions being significantly higher than 0.5, except I^+^R^0^ (Table 2).

**Table 2.** Mean static Bayesian model priors vs 0.5 for blocks 2-4.

| Condition | Mean prior | $t$ | $p$ | $\text{BF}_{10}$ |
| --- | --- | --- | --- | --- |
| $I^0R^+$ | 0.53 | 2.7 | 0.007 | 3.5 |
| $I^+R^0$ | 0.50 | 0.5 | 0.64 | 0.12 |
| $I^+R^+$ | 0.63 | 7.3 | $10^{-9}$ | $10^6$ |
| $I^-R^+$ | 0.61 | 5.5 | $10^{-6}$ | $10^4$ |

*The static Bayesian model gives one prior value for each participant. We compare these to the uninformed prior of 0.5 using two-sample t-tests. Priors for all conditions except $I^+R^0$ were significantly higher than 0.5.*

**Figure 4.**
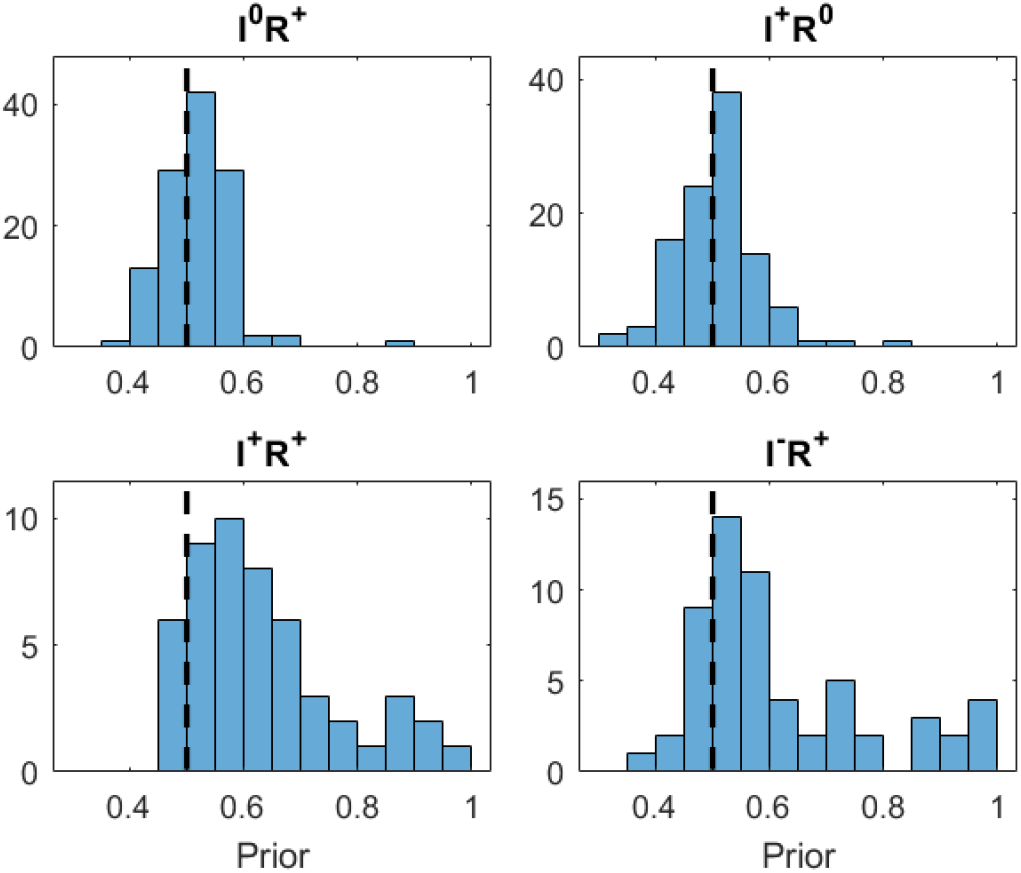
Histograms of prior values for all conditions. The dashed line shows the neutral prior of 0.5.

### Autistic traits

We investigated the relationship between autistic traits and both behavioural and model-based measures. Contrary to our hypothesis, autistic traits were not correlated with biases (F = 1.0, *p* = 0.32, BF_10_ = 10^−6^), model-estimated priors (F = 0.7, *p* = 0.39, BF_10_ = 10^−6^, Figure 5), or model-estimated likelihoods (F = 1.0, *p* = 0.31, BF_10_ = 10^−6^). Nor was there any interaction between conditions and AQ scores in how they influenced biases (F = 0.2, *p* = 0.88, BF_10_ = 10^−4^), priors (F = 0.38, *p* = 0.77, BF_10_ = 10^−4^), or likelihoods (F = 0.1, *p* = 0.98, BF_10_ = 10^−4^). To investigate the possibility that autistic traits influenced the acquisition of the regularities, we also fit our static Bayes model exclusively to data from the first block, but found no interaction between AQ scores and conditions (F = 2.0, *p* = 0.12, BF_10_ = 0.004).

**Figure 5.**
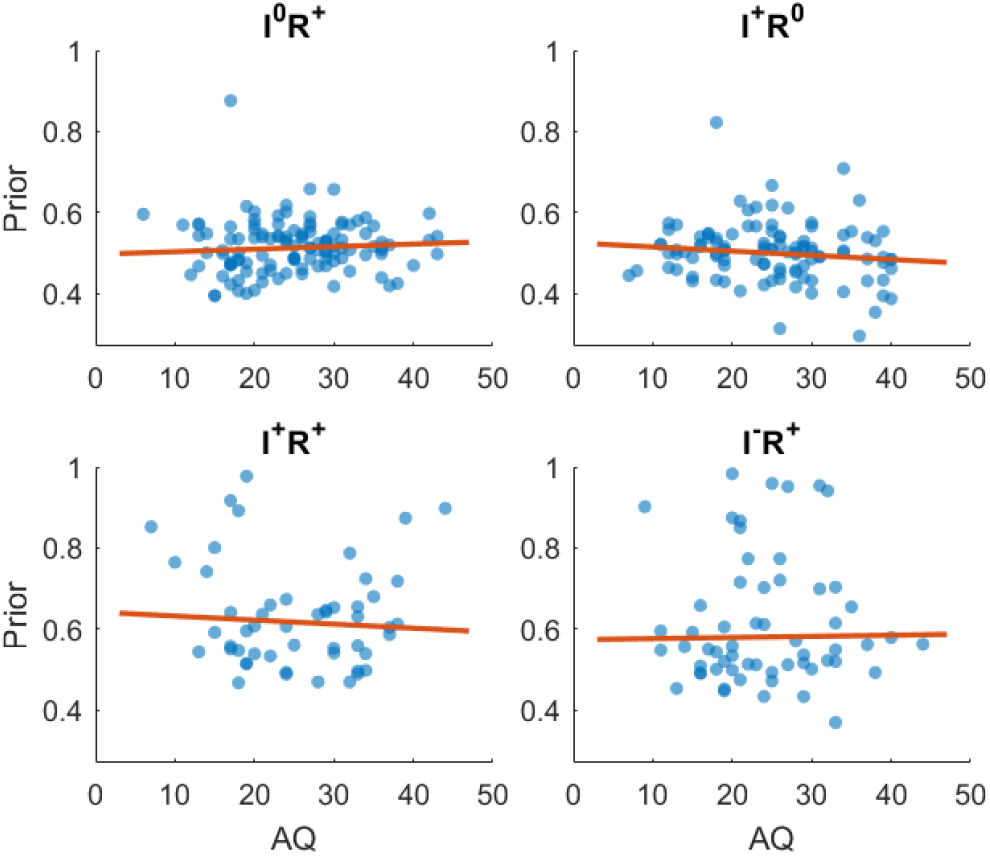
AQ scores vs participant priors. The results show no relationship in any condition.

## Discussion

### Implicit and explicit learning

In this study, we explored how the learning of probabilistic associations between auditory cues and visual stimuli is impacted by the presence and the content of explicit instructions and how it is affected by autistic traits. We designed four experimental conditions that differed in the presence of associations and the veracity of the instructions given to participants. We expected participants to initially form a Bayesian prior based on the instructions of the task, which would then be updated following the statistics of the stimuli. Indeed, our results showed that participants were influenced by both the information explicitly provided to them and the regularities of the task.

Specifically, when participants were not alerted to the presence of any association between the cue and stimulus, they rapidly formed priors in line with the environmental regularities. However, through Bayesian modelling, we estimated these priors to be significantly weaker than the actual regularities (53% vs 75%), although they were still significantly different from chance (50%). In another condition, where participants were falsely informed about the presence of an association, they initially showed a noticeable bias that diminished rapidly as they progressed in the task. Remarkably, in conditions where implicit and explicit information coexisted, participants developed more robust priors in the direction of the regularities (61%-63%), independently of the veracity of the instructions. This comes in contrast to our expectations, with misleading instructions having a positive effect on the participants’ learning of the regularities. It also contrasts with previous work which has shown that searching for regularities inhibits the learning of environmental statistics [7,14,15].

Compared to previous studies, one difference with our task is that past work has mostly used serial reaction time and artificial grammar learning tasks. In both of these, the regularities that participants had to learn were rule-based, as opposed to our own task where regularities were frequency-based. For example, in some of these tasks, there would be a numeric sequence repeating (e.g., 1-1-3-1-4-2-2-4-1) and participants would have to respond as quickly as possible to each number by pressing the assorted key. In others, participants might have to learn that words with specific substrings are valid (e.g., VSXS and XXV) and others are not. In both of them, the regularities of the environment are far more precise (akin to rules) than a simple difference in the frequence of some stimuli or some probabilistic association between them. One would naively expect that being aware of the presence of regularities and consciously searching for them (explicit learning) is better suited for precise rules, while unconscious or implicit learning is better suited for general statistical patterns. However, this is the opposite pattern of what the results of this and past studies suggest. One possible explanation for the observed detrimental effect of instructions in the rule-based tasks could be that their exact regularities are too complicated to be learned during a single experiment. At the same time, implicit learning might be able to extract some useful patterns, even if these are far from the exact rules. In the aforementioned example sequence, implicit learning might result in a general prior of the form ‘3s and 1s go together, 2s and 4s go together’. This obviously cannot explain the full sequence, but it is nonetheless useful. In contrast, participants who explicitly search for the exact form for the regularities might disregard incomplete rules, while dedicating many resources to discovering the exact pattern, leading to worse performance and no discernible benefits. In our tasks, on the other hand, explicit instructions would be perfectly able to provide an understanding of the true regularities, especially given that instructions explained the general nature of these patterns, if not their actual form.

### Autistic traits

We investigated the relationship between autistic traits and task behaviour. Our hypothesis was that autism would be associated with weaker biases, primarily in the implicit condition. This was refuted by our results, as they showed no relationship between autistic traits and the magnitude of Bayesian priors or likelihoods across conditions and no interaction between the conditions and AQ scores. They also showed no effect of AQ during prior acquisition.

Our literature review [21] had revealed that reduced biases in autism were most commonly observed in implicit learning tasks, while explicit information appeared to help compensate for prior acquisition deficits. However, the findings were inconsistent – some implicit learning studies showed intact priors, while some explicit learning studies found differences between groups. We proposed several explanations for these mixed results. First, task classification could have been problematic since instructions are often omitted from published articles, making it difficult to definitively categorise tasks as implicit or explicit. Second, we encountered various types of explicit priors: participants being falsely told about the presence of regularities [25], receiving accurate information about the actual regularities [26], or simply being informed that regularities exist without specific details [19]. Lastly, the review revealed considerable experiment heterogeneity, with no single task being used to test both implicit and explicit prior acquisition.

Our experiment was designed to address all of these issues. We used the exact same task across all conditions, and we included both a condition where participants are given true information about the regularities as well as conditions where participants are provided with false information. Moreover, the instructions in our task had a strong effect on participant behaviour, showing that implicit and explicit conditions could be clearly differentiated. Despite that we found no influence of autistic traits in any of the conditions.

On top of our investigations using the static Bayesian model, we also performed some exploratory analysis of the belief trajectories of the participants using the Hierarchical Gaussian Filter (Section 6 in S1 Supplementary Information). This revealed a weak correlation between autistic traits and learning rates in the implicit condition, but not in the other conditions. Given that the HGF was not the winning model, we can draw no conclusions from this result. However, it could constitute a fruitful direction of future research for both the theory of volatility overestimation [19,20] and the theory of weaker priors [17] (see Section 6 in supplementary information for further discussion).

### Conclusions

Our study demonstrates how explicit information might influence probabilistic learning, with participants’ prior acquisition being significantly strengthened by instructions, even when those were inaccurate. This comes in contrast with previous results from rule-based tasks, potentially due to the frequency-based nature of our task. Our findings emphasise that instructions are not merely procedural details but experimental variables that can fundamentally alter participant behaviour. Moving forward, studies should specify the type of learning they aim to evoke and document instructions with the same level of detail as other methodological parameters. Surprisingly, our results did not demonstrate any differences in prior strength across autistic traits in either the implicit or the explicit conditions, contrary to our hypothesis and the most prominent Bayesian theories for autism.

