## Supplementaray information for "Influence of truthful and misleading instructions on statistical learning across the autism spectrum"

### S1 Supplementary Information

Nikitas Angeletos Chrysaitis & Peggy Seriès

### 1. Sample characteristics and AQ scores

The recruiting platform we used, Prolific, asks the participants various questions. During recruiting, we used on if these, 'Have you received a formal clinical diagnosis of autistic spectrum disorder, made by a psychiatrist, psychologist, or other qualified medical specialist? This includes Asperger's syndrome, Autistic Disorder, High Functioning Autism or Pervasive Developmental Disorder', to ensure that we would get a broad enough range for autistic traits. The question had the following possible answers:

1. Yes – as a child
2. Yes – as an adult
3. I am in the process of receiving a diagnosis
4. No – but I identify as being on the autistic spectrum
5. No
6. Don't know / rather not say

We recruited half of our Prolific participants from those that had chosen options 1-4, and half from those that had chosen option 5. Specifically in our final sample, the percentage of participants that had chosen option 5 was 50% for $I^0R^+$, 48% for $I^+R^0$, 41% for $I^+R^+$, and 44% for $I^+R^0$. AQ scores did not differ significantly across conditions (one-way ANOVA: F = 0.3, $p$ = 0.83, $BF_{10}$ = 0.006). The conditions did not significantly differ in the gender or age distribution (chi-squared test: $\chi^2$ = 2.5, $p$ = 0.87; one-way ANOVA: F = 2.1, $p$ = 0.11, $BF_{10}$ = 0.06).

To measure the reliability of the AQ scores, we used the split-half correlation. The questionnaire was split in half at random and we calculated 2 AQ scores based on each half.

Then we measured the Pearson correlation between them. Because the calculated correlation was based only on half the questionnaire, we adjusted it using the Spearman-Brown formula, with the adjusted correlation being r_adj = 2·r_half / (1 + r_half). This process was repeated for 10000 random splits of the dataset. The average correlation was 0.85, indicating good reliability of the AQ scores.

### 2. Stimulus Contrast

Running the experiment online means that participants take part under very different lighting conditions. To ameliorate that, we included a contrast staircase in our experiment. The staircase was set to change rapidly during the training trials, so that it would adjust quickly to the participants' conditions, and then to become more stable during the main experiment. We tried to have the staircase reach a low enough value in the training so that biases would be present during the first trials of the experiment. If the contrast was set too high, the participants would have plenty of information from the gratings and would ignore the auditory cue. This had the unintended side-effect that the contrast was set too low and slightly adjusted during the experiment at a rate of 7%-16% (Figure 1). Interestingly, this happened predominantly in the $I^+R^0$ and $I^-R^+$ conditions, in which misleading information was given to the participants.

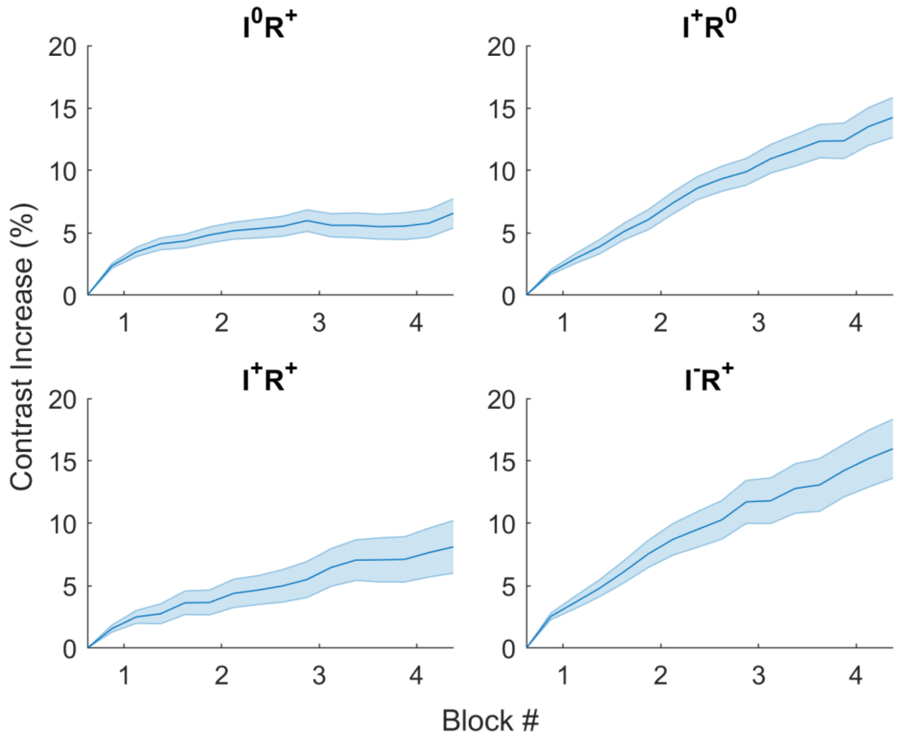


*Figure 1: Contrast change during the main experiment.*

### 3. Details of the learning models

In the HGF the first level ($x_1$) represents the outcome of a specific trial (left- or right-tilted grating), the second one ($x_2$) represents the association between the tones and the gratings over time, and the third one ($x_3$) the volatility of the environment, i.e. the rate of change of the association. Mathematically, we have

$$x_1^{(t)} \sim \text{Bernoulli}\left(s\left(k_1 x_2^{(t)}\right)\right)$$

$$x_2^{(t)} \sim N\left(x_2^{(t-1)}, \exp\left(k_2 x_3^{(t)} + \omega_2\right)\right)$$

$$x_3^{(t)} \sim N\left(x_3^{(t)}, \exp(\omega_3)\right)$$

with $s(x) = 1 / (1 + \exp(-x))$. The output of the HGF is the prior for $x_1$ at time $t$, called $Pp$ in the main text.

The model has the following free parameters $k_1$, $k_2$, $\omega_2$, $\omega_3$, $\mu^0{}_2$ and $\sigma^0{}_2$ (which defined the distribution for $x_2$ at $t = 0$), $\mu^0{}_3$ and $\sigma^0{}_3$ (which defined the distribution for $x_3$ at $t = 0$), and $c_0$ (from the definition of $Pv$).

In the Rescorla-Wagner model, the prior starts at value $Pp^0$ and is dynamically adjusted based on the prediction error between that and the stimulus:

$$Pp^t = Pp^{t-1} + a \cdot (u^t - Pp^{t-1})$$

with $Pp^0$, $a$, and $c_0$ being the free parameters.

### 4. Model Comparison

We compared a great variety of HGF variants using Bayesian model comparison in all four conditions of our task and used the best performing version for the comparison with the other models. In all conditions, the best HGF assumed that the environment stayed stable over time. That is, $x_3$ was set at $-\infty$, with $k_2$, $\omega_2$, $\omega_3$, $\mu^0{}_3$, and $\sigma^0{}_3$ being fixed, and the change in second-level beliefs resulted only from acquiring more information from the environment. The only difference between the models was the starting point of the second-level beliefs, which was set at 0 in the $I^0R^+$ condition. All of the winning models also had $k_1$ as a free parameter, which mediates the relationship between second and first-level beliefs.

***Table 1. Posterior model probabilities.***

| | | Models | | | |
|---|---|---|---|---|---|
| | | response model | static Bayesian | HGF | RW |
| Conditions | $I^0R^+$ | 0.03 | 0.95 | 0.02 | 0.01 |
| | $I^+R^0$ | 0.04 | 0.82 | 0.08 | 0.05 |
| | $I^+R^+$ | 0.09 | 0.85 | 0.02 | 0.04 |
| | $I^-R^+$ | 0.09 | 0.77 | 0.02 | 0.12 |

### 5. Model & Parameter Identifiability

To ensure that the results of Table 1 were reliable, we performed model recovery using the same models. We simulated 500 participants using each model and then fitted them with all candidate models. We then looked at the percentage of simulated participants that were best fit by each model. Ideally a model would fit best 100% of the participants that it simulated and 0% of those that the other models simulated. Our results showed medium to poor identifiability for all models besides the Rescorla-Wagner, as many of the simulated participants were recovered by other models than the one that had simulated them across conditions. The HGF specifically was poorly recovered in all conditions. This could be due to the fact that most participants learned the regularities very quickly in our task and therefore only a few trials would show a changing prior.

***Table 2. Model recovery.*** *The table shows the percentage of simulated participants that were recovered by each model.*

| | Recovered Participants | | | | |
|---|---|---|---|---|---|
| | **$I^0R^+$** | response model | static Bayesian | HGF | RW |
| | response model | **44** | 33 | 21 | 1 |
| | static Bayesian | 41 | **48** | 11 | 0 |
| | HGF | **37** | 35 | 26 | 1 |
| | RW | 0 | 0 | 1 | **99** |
| | **$I^+R^0$** | response model | static Bayesian | HGF | RW |
| | response model | **53** | 43 | 3 | 1 |
| | static Bayesian | 35 | **45** | 10 | 1 |
| | HGF | 31 | **42** | 23 | 5 |
| Simulated Participants | RW | 0 | 2 | 0 | **98** |
| | **$I^+R^+$** | response model | static Bayesian | HGF | RW |
| | response model | **39** | 37 | 22 | 2 |
| | static Bayesian | 36 | **50** | 12 | 1 |
| | HGF | 32 | **40** | 25 | 4 |
| | RW | 0 | 0 | 0 | **100** |
| | **$I^-R^+$** | response model | static Bayesian | HGF | RW |
| | response model | **48** | 39 | 12 | 0 |
| | static Bayesian | 33 | **52** | 14 | 1 |
| | HGF | 23 | **41** | 29 | 6 |
| | RW | 0 | 1 | 0 | **99** |

We also performed parameter recovery, by simulating the responses of 1000 participants with our winning, static Bayesian model and then refitting them using the same model. Pearson correlations between the simulated and recovered parameters showed good identifiability for both the prior and the baseline level of contrast which determines the likelihood values.

***Table 3. Parameter recovery.*** *Pearson correlations between simulated and recovered parameter values. mu_0 is the starting point of the second-level beliefs, sa_0 is the uncertainty at that point.*

| | | Parameters | |
|---|---|---|---|
| | | *Pp* | $c_0$ |
| Conditions | $I^0R^+$ | 0.69 | 0.86 |
| | $I^+R^0$ | 0.75 | 0.90 |
| | $I^+R^+$ | 0.87 | 0.79 |
| | $I^-R^+$ | 0.90 | 0.88 |

### 6. Exploratory analysis using the Hierarchical Gaussian Filter

As our task involved learning the environmental regularities, we decided to also model participant behaviour using the HGF, despite it not winning our model comparison. The model recovery results (Table 2) demonstrate that even if the cognitive processes underlying participant behaviour were better approximated by the HGF, our model comparison might be unable to reflect that.

We analysed the trajectories of HGF-estimated first-level priors used by the participants ($x_1$). Participants initially formed a belief based on the instructions, as demonstrated by their prior at the first trial being significantly different from the uniform prior 0.5 ($I^+R^0$: M = 0.61, $t$ = 4.3, $p = 10^{-5}$; $I^+R^+$: M = 0.57, $t$ = 2.7, $p$ = 0.01; $I^-R^+$: M = 0.44, $t$ = –2.0, $p$ = 0.052). But when that belief disagreed with the actual regularities, it was quickly amended, with no difference between the priors of $I^+R^+$ and $I^-R^+$ after the first block ($t$ = 1.1, $p$ = 0.25, $BF_{10}$ = 0.36). Priors in the $I^+R^+$ and the $I^-R^+$ conditions were much stronger than priors in the implicit condition ($t$ = 7.9, $p = 10^{-13}$, $BF_{10} = 10^7$ and $t$ = 5.2, = $10^{-6}$, $BF_{10}$ = 1598).

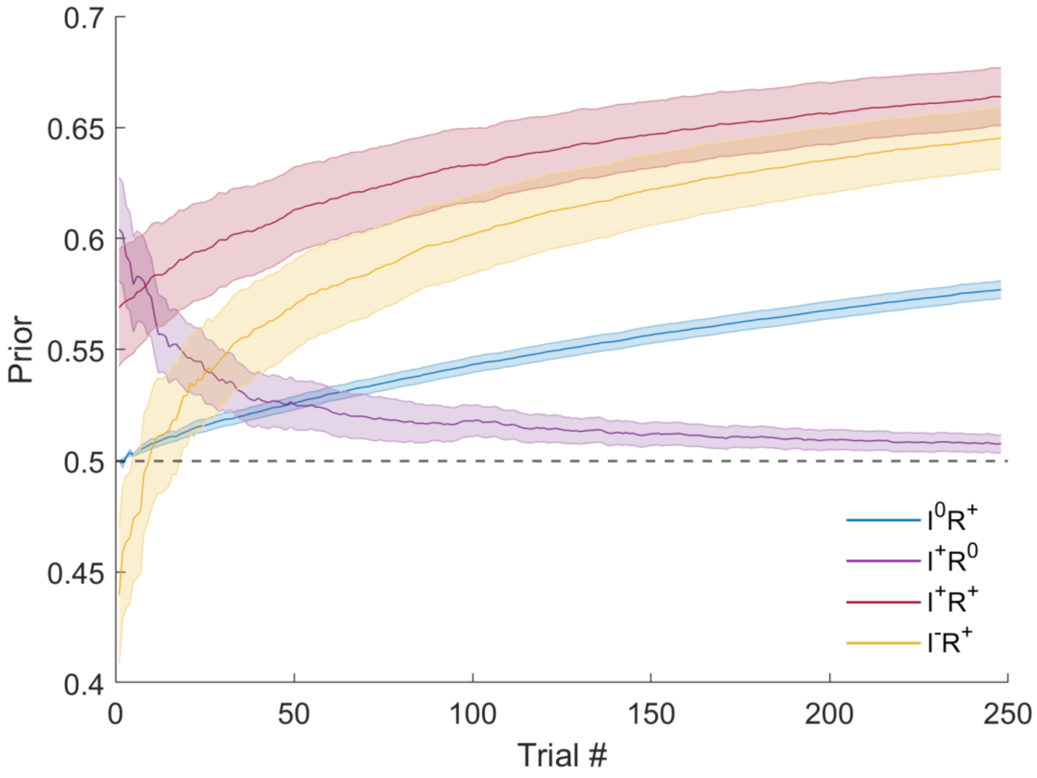


***Figure 2: The trajectories of model-estimated first-level priors in the four conditions.*** *Priors were significantly affected by the instructions in the beginning of the task, but that was quickly amended based on the regularities in each condition. Priors in the conditions that combined explicit information and regularities were much stronger than those in the implicit condition.*

One difference between these and the behavioural findings is that the computationally estimated first-level prior in the $I^+R^0$ condition never reached 0. Moreover, while the biases in Figure 3 of the main text remained relatively stable after the first block, Figure 2 in the Supplementary Information shows the priors increasing in three out of four conditions even in the final trials. On top of these, the estimated priors start at different points among the $I^+R^+$, and $I^-R^+$ conditions, despite the instructions being identical. We suspect that these effects are artifacts cause by the structure of the HGF and they do not reflect the participants' cognitive processes.

We looked at Pearson correlations between HGF behaviour and AQ scores. To avoid outliers disproportionately influencing the results, we winsorised the data, meaning that we equated the outliers (determined using the interquartile method) with the most extreme non-outlier values [1]. This allowed us to keep some of the information in the outliers, while mitigating their outsized influence. We found no relationship between AQ and model-estimated first-level priors (F = 0.44, $p$ = 0.51, $BF_{10} = 10^{-6}$), nor an interaction between AQ and conditions (F = 0.25, $p$ = 0.86, $BF_{10} = 10^{-4}$). We also investigated the relationship between AQ scores and learning rates or prior variances (Figure 3). The resulting Pearson correlations showed a marginal effect in the $I^+R^0$ condition ($r = -0.19$, $p = 0.06$) and a significant positive correlation in the $I^0R^+$ condition ($r = 0.19$, $p = 0.03$), that however would not survive a correction for multiple comparisons.

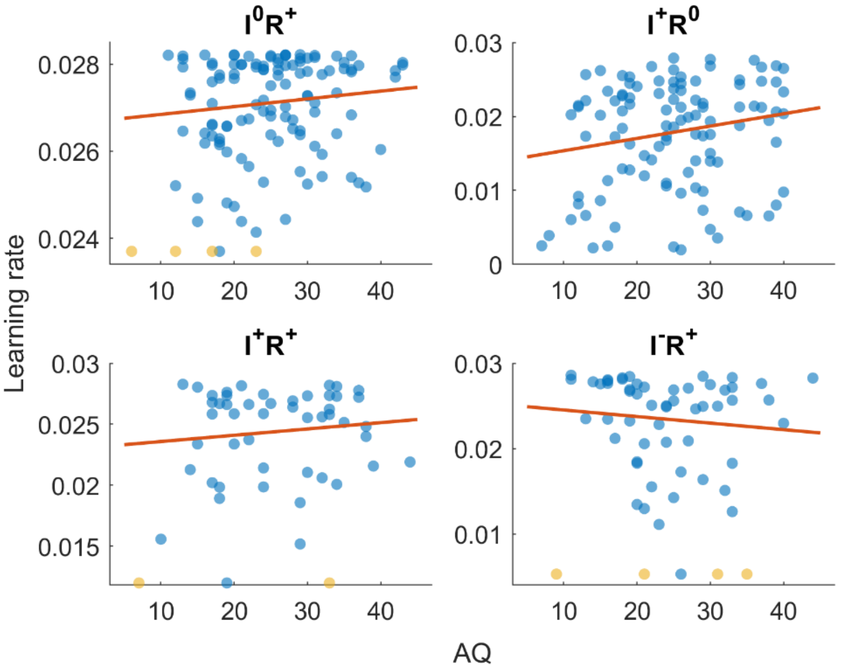


***Figure 3: Correlations between AQ and average second-level learning rates (variances).*** *Orange dots correspond to winsorised outliers, and the red line to a robust linear regression fit. In the implicit condition the correlation was r = 0.19, p = 0.03.*

### 7. Discussion of the exploratory results

The presence of a correlation between AQ scores and second-level uncertainty might appear contradicting to the absence of a negative correlation between AQ and participant biases (main results section Autistic Traits), but this is not necessarily the case. In tasks such as the one presented in this paper, prior beliefs about individual trials do result from beliefs about task regularities but are distinct from them. Individual trial priors can be represented by a single number: after the tone is heard, how likely is it that the grating will be tilted to the left? Priors closer to 0 or 1 correspond to large certainty about what the stimulus would be (left or right). Priors at 0.5 are fully uncertain, not biasing responses towards any direction and completely amenable to new information. Prior beliefs about the task regularities, on the other hand, cannot be represented in the same way. These beliefs consist of *distributions* over these probabilities. In that case, a strong, certain prior is one that is more concentrated around a specific value, independently of what that value is. A completely uncertain prior would be one that is uniform over all possible values. Take the examples in Figure 4. There we see a strong prior that the experiment has no regularities and a weak prior that there is a 65% association between cue and stimulus. Despite the latter being a less certain belief about the regularities, it would result in a *stronger* prior for individual trials. This is because the precision of beliefs about regularities determines how these priors are updated, not their influence during perception. To give another example, a strong belief that a coin is fair would result in a very weak belief about the coin landing heads or tails in an individual flip.

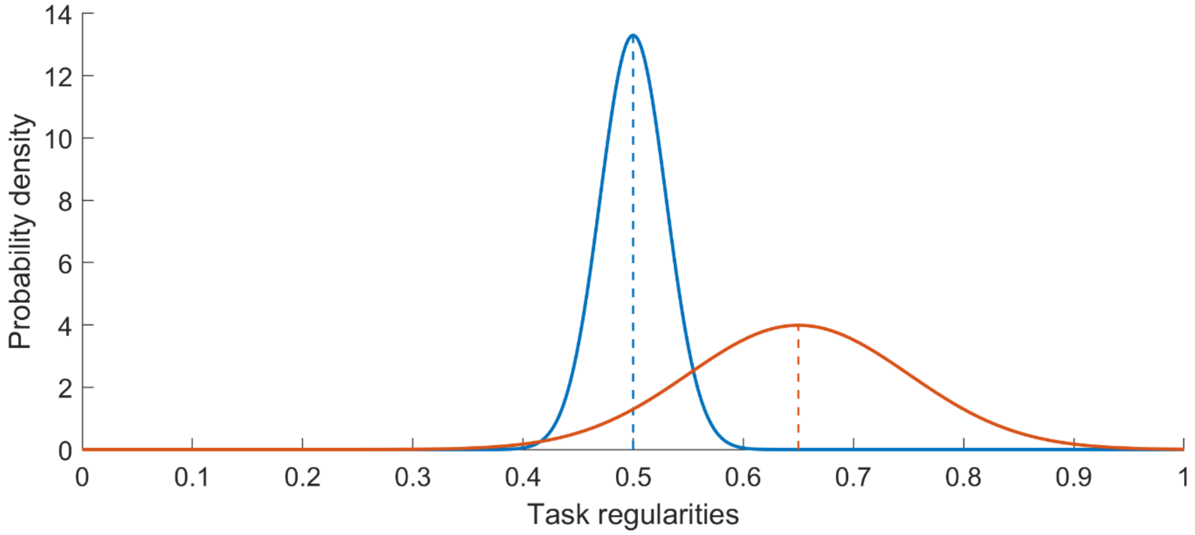


***Figure 4: Examples of beliefs about task regularities.*** *The blue line corresponds a strong belief around 0.5, the red line corresponds to a weak belief around 0.65. Dashed lines show the mean of the distributions. In the Hierarchical Gaussian Filter these priors are encoded in the second-level of the model and their mean is used to form the first-level prior for the individual trials.*

In the HGF priors about individual trials are represented by the first level of the model, while prior beliefs about regularities are represented by the second level. Our results showed no correlation between autistic traits and first-level priors or the corresponding response biases, but could constitute weak evidence for more uncertain second-level priors. This means that the theory of Pellicano and Burr [2] could be correct and not produce weaker biases in autism. Moreover, higher learning rates would be predicted from the theory of Palmer et al. [3] and specifically its instantiation that hypothesises volatility overestimation in autism [4].

Building on that, our results showed that this relationship goes away when participants are alerted about the presence of regularities. Therefore, it is possible that individuals with autism use explicit strategies to compensate for implicit learning differences. In the seminal paper by Lawson et al. [4], participants were informed about the presence of regularities and that these would change over time. Their results showed no relationship between diagnoses and the second-level volatility estimates of the participants ($\omega_2$). However, participants were not instructed that the rate of change would itself change during the task. And indeed the authors show differences across diagnostic groups in how the learning rates changed during the task and higher third-level volatility estimates ($\omega_3$) in ASD.
